# Duplex formation between the template and the nascent strand in the transcription-regulating sequences determines the site of template switching in SARS – CoV-2

**DOI:** 10.1101/2020.12.11.416818

**Authors:** Aaron R. D’souza, Amanda Buckingham, Fanny Salasc, Carin Ingemarsdotter, Gennaro Iaconis, Isobel Jarvis, Harriet C. T. Groom, Julia Kenyon, Andrew M. L. Lever

## Abstract

Recently published transcriptomic data of the SARS-CoV-2 coronavirus show that there is a large variation in the frequency and steady state levels of subgenomic mRNA sequences. This variation is derived from discontinuous subgenomic RNA synthesis where the polymerase switches template from a 3’ proximal genome body sequence to a 5’ untranslated leader sequence. This leads to a fusion between the common 5’ leader sequence and a 3’ proximal body sequence in the RNA product. This process revolves around a common core sequence (CS) that is present at both the template sites that make up the fusion junction. Base-pairing between the leader CS and the nascent complementary minus strand body CS, and flanking regions (together called the transcription regulating sequence, TRS) is vital for this template switching event. However, various factors can influence the site of template switching within the same TRS duplex. Here, we model the duplexes formed between the leader and complementary body TRS regions, hypothesising the role of the stability of the TRS duplex in determining the major sites of template switching for the most abundant mRNAs. We indicate that the stability of secondary structures and the speed of transcription play key roles in determining the probability of template switching in the production of subgenomic RNAs.

## INTRODUCTION

The human coronaviruses include a diversity of viruses that cause diseases such as the common cold as well as those resulting in epidemics and pandemics, such as the Middle East Respiratory Syndrome coronavirus (MERS-CoV), Severe Acute Respiratory Syndrome coronavirus (SARS-CoV) and SARS-CoV-2. They belong to the family Coronaviridae in the order Nidovirales ^1,2^. Coronavirus genomes consist of a positive sense single stranded RNA molecule of approximately 30 kb in length. They share a common architecture that includes highly structured 5’ and 3’ terminal untranslated regions (UTRs) flanking two overlapping non-structural polyprotein encoding open reading frames 1a and 1b (ORF1a, ORF1b). 1a/1b comprises the 5’ two-thirds of the intervening sequence and the remaining one-third of the genome encodes structural and accessory proteins. The structural proteins include the spike protein (S), envelope protein (E), membrane protein (M) and nucleocapsid protein (N). The six accessory proteins present in SARS-CoV-2 include 3a, 6,7a,7b, 8 and, putatively, 10 ^3^.

The replication of the genome and the production of mRNAs for translation are intimately linked processes. They both require RNA-dependent RNA synthesis, beginning with the synthesis of a negative sense strand from the incoming positive sense genome by the virus-encoded RNA-dependent RNA polymerase. This begins at the 3’ terminus of the positive strand and moves in the (negative sense) 5’ – 3’ direction ^4^. Full length and truncated negative strands are then used as templates to amplify positive strand RNAs. The full genome length RNA (gRNA), containing ORF1a and ORF1b, is used for translation and/or packaging, while smaller positive strand subgenomic mRNAs (sgRNAs), encoding structural and accessory proteins, are primarily destined for translation ^4,5^. In SARS-CoV-2 infected cells, there are nine canonical positive sense sgRNAs that are produced in addition to the gRNA, each identified by the product of their 5’-terminal open reading frame – S, 3a, E, M, 6, 7a, 7b, 8 and N.

The canonical sgRNAs are produced through a process called discontinuous transcription, conserved in both arteriviruses and coronaviruses ^6^. These positive strand mRNAs contain a common 5’ leader sequence (70 - 100 nt) derived from the 5’ terminus of the positive strand genome ^7^. During canonical discontinuous transcription, chimeric mRNA molecules are produced each consisting of an identical segment of the leader sequence fused with one of several non-contiguous regions in the body of the genome found 5’ of each ORF. Pseudo-circularisation of the genome by unknown viral and/or cellular factors brings together transcription-regulating sequences (TRS) found in the leader (TRS-L, 5’) and at the 5’ end of the body sequence (TRS-B, 3’) ^1^. The TRS includes a conserved core sequence (CS) of 6-7 nt common to both the TRS-L and the TRS-B ^8^; in SARS-CoV-2 this is ACGAAC. Thus, a duplex is formed between the CS-L and the complementary CS-B (cCS-B) in the nascent negative strand leading to a switch in the polymerase template from the body sequence to the 5’ leader sequence. If this does not occur, the polymerase progresses until it either encounters a subsequent TRS and makes a template switch to the 5’ positive strand leader or completes the full antigenomic strand (van Marle et al. 1999; Pasternak et al. 2001; Zúñiga et al. 2004).

The 5’ UTR consists of a series of stem-loops (SL), not all of which are conserved between different coronaviruses. SL1 and SL2 are highly conserved and are required for genome replication and possibly RNA production ^12–14^. The SL3 stem-loop, present in SARS-CoV-2, is not universally conserved but this region contains sequences important for transcription in all coronaviruses ^15^. Downstream of this is a conserved SL4 hairpin that may be responsible for orientating the SL1, SL2 and SL3 regions during transcription ^16^. The 5’ UTR ends with a large SL5 stem-loop consisting of a 4-way junction. It is common to many SARS-related viruses and contains of the ORF1ab start codon. It has been proposed to function as a packaging signal 17.

The role of the various conserved stem-loops of the 5’ UTR in discontinuous transcription is unclear. SL1, SL2 and SL4 show functional conservation between viruses; swapping them between related coronaviruses yields viable chimeric viruses unless the TRS is also replaced ^14,18,19^. In mouse hepatitis virus (MHV), mutational analyses showed that neither the sequence nor structure of SL4 is vital for viral replication. However, deletion of SL4 was found to inhibit sgRNA production ^14,16^. Experiments involving mutation or deletion of part of, or the complete stem-loop suggest that the structural features of these stem-loops are more important than their nucleotide sequence ^1,19^. However, their exact function remains unclear.

Recent modelling informed by RNA structure probing generated a single nucleotide resolution map of the secondary structures in the SARS-CoV-2 genome. This predicted a high degree of structure compared to other positive sense RNA viral genomes such as those of HIV-1 and ZIKV ^20^ and even more structure than the related viruses, MERS and SARS-CoV ^21^. The probing confirmed that an estimated 38.6% of the SARS-CoV-2 genome is structured with around 8% of the structures exhibiting significant covariance with other coronaviruses, implying that these structures are conserved between the different viruses and thus of functional importance (Manfredonia et al. 2020). These experiments confirmed structure and sequence conservation in the SL1-SL4 stem-loops between many coronaviruses. Unlike the corresponding structure of some other coronaviruses, the SARS-CoV-2 ACGAAC CS-L is found in the SL3 stem-loop with part of the sequence exposed in an apical loop. A similar architecture has been described in some other coronaviruses where the CS is also found in a stem-loop ^17,20,23^.

The stability of the TRS duplex is a key determinant of the occurrence of and rates of template switching. Mutations in the TRS-L of Equine arteritis virus (EAV) exposed new sites of complementarity in the genome leading to the production of novel minor sgRNAs ^24^. Furthermore, changing the level of complementarity by mutating the TRS-L or changing the length of the complementary TRS-flanking regions led to a higher or lower incidence of template switching depending on whether the stability of the duplex formed between the TRS-L in the positive strand and the complementary sequence in the nascent negative strand is increased or decreased respectively ^24^.

If the canonical TRS-L is mutated, alternate cryptic TRSs, downstream of the site of the canonical TRS and upstream of the initiation codon, can be recruited which allow production of sufficient amounts of sgRNAs to yield infectious progeny virus albeit at a reduced titer ^25^. By contrast, the transmissible gastroenteritis virus (TGEV) genome contains a second canonical CS (CS-2) in its S gene that does not initiate discontinuous transcription and the production of detectable levels of mRNA. Base pairing between the CS-2 and complementary leader sequence alone is seemingly not sufficient to cause template switching. In many cases, base pairing of adjacent nucleotides flanking the CS is vital for polymerase strand transfer. This phenomenon is genome position-independent ^26^ and lends credence to a model in which template switching is primarily determined by the overall free energy of the duplex formed and not specifically defined by the core sequence or by its location in the genome.

Transcriptomic analysis of members of the order Nidovirales has identified multiple non-canonical fusion events resulting in additional sgRNAs to those produced by template switching at cTRS-B/TRS-L sequences. Some of these lead to translation products derived from alternative open reading frames identified by mass spectrometry and ribosome profiling ^27^. Recent transcriptomic analysis of SARS-CoV-2 showed that in addition to the canonical TRS-mediated leader-body fusion events, which account for ~93% of the transcripts, recombination events also occur between regions where only one or neither of the two TRSs is present. Moreover, there were a variety of leader-body junction sequences associated with each canonical ORF, suggesting that there are additional factors that govern template switching ^3^. Here, we use published transcriptomic data to model the cTRS-L:TRS-B duplex structures proposed by the discontinuous transcription model and use this to infer how duplex formation and secondary structure determine the probability of template switching.

## METHODS

The leader-body junction sequences were extracted from the supplementary data provided in Kim et al. (2020). Since all the template switching events that occurred within the duplex or immediately after duplexes would produce identical fusion junctions, the region of possible template switching was identified and a sequence was constructed in silico containing 94 nucleotides of the 5’ UTR (uppercase), a 10 nt ‘spacer’ that is forced to be single-stranded and a nascent negative strand sequence (uppercase) of varying lengths ending at the last possible site of the template switching or up to 6 nucleotides after this position (Supplementary figure 1). Then, RNA secondary structure was modelled using the RNAstructure software ^28^. Summarised structures were drawn using the XRNA software ^29^. Sites of template switching were marked using information provided by Kim et al. (2020).

## RESULTS

SARS-CoV-2 transcriptomic data published by Kim et al., (2020) showed that the most abundant sgRNAs derive from fusion events involving both the TRS-L and one or other TRS-B sequence. The top 24 sgRNAs analysed in this paper comprise those products of discontinuous transcription that are above 0.01% of the total transcripts and are mostly derived from a TRS-L:cTRS-B fusion event. Despite sharing common core sequences, there are large differences in the abundance of the individual sgRNAs. The reasons for this are not obvious but point to there being multiple factors controlling the rates of template switching and the quantity of individual sgRNAs generated. Moreover, multiple different variants of the same sgRNA were found that had utilised different sites of template switching. Examining the relative abundance of the canonical and non-canonical sgRNAs, of the nine canonical sgRNAs (S, 3a, E, M, 6, 7a, 7b, 8, N), ORF7b is fourteenth in the order of most abundant sgRNAs produced (Fig. 1). By contrast four different versions of the sgRNA encoding the nucleocapsid (N) protein appear before the most abundant ORF7b sgRNA. Plausibly this may be due to the non-canonical core sequence (AAGAAC) that occurs in the ORF7b TRS-B. There are also non-canonical sgRNAs that are produced where the body junction is found within the ORF1ab open reading frame (labelled ORF1ab, producing a putative 28 aa N-terminal peptide) or within the N open reading frame (labelled N*, producing a putative 103 aa peptide) (Fig 1).

**FIGURE 1.**
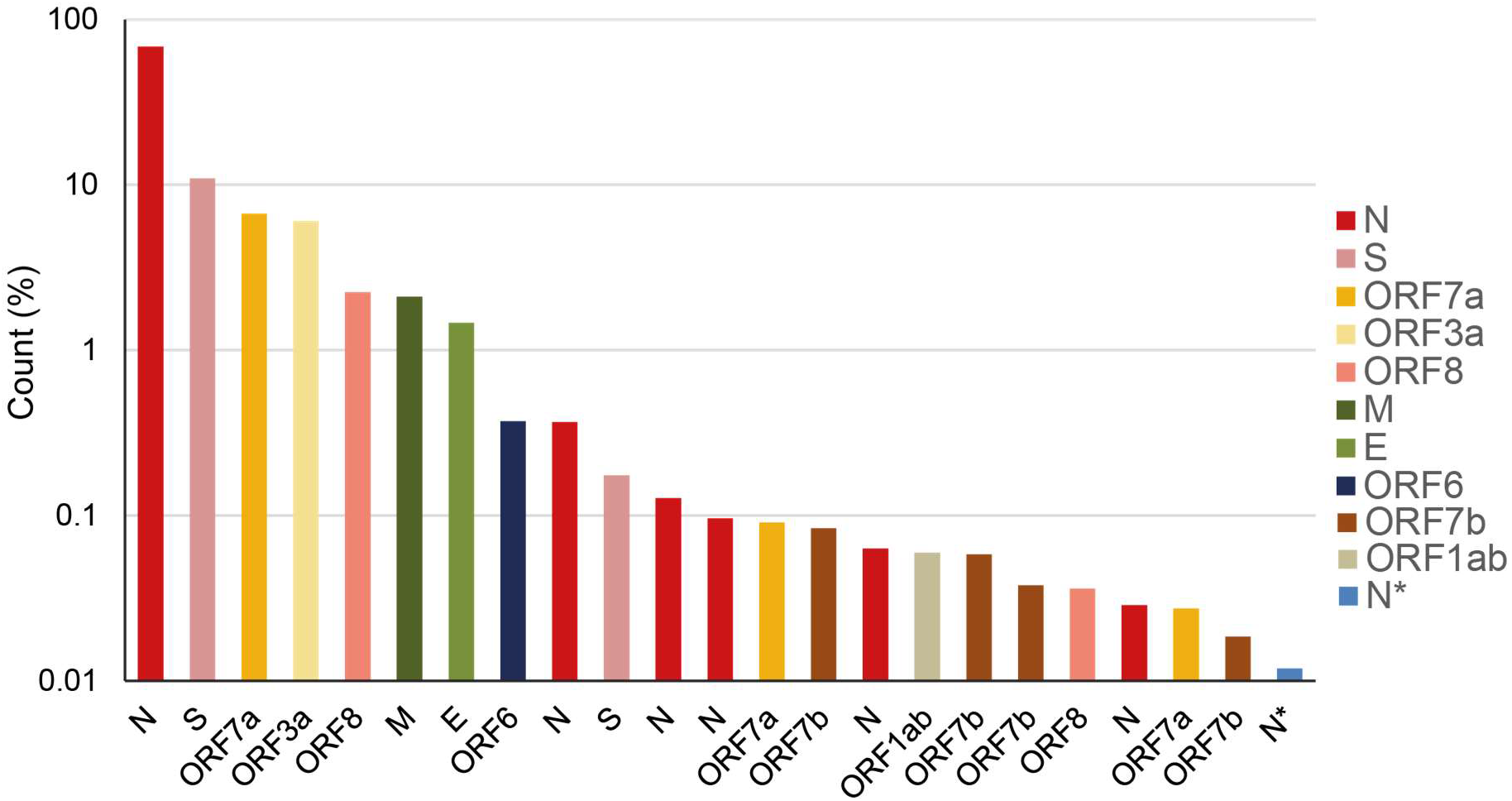
Percentage abundance of the major transcripts above 0.01% of the total transcript abundance from the DNA Nanoball Sequencing data published by Kim et al. (2020). These transcripts were produced from discontinuous transcription events from the body of the genome to the 5’ leader. The key indicates the location of the body sequence junction point.

### The TRS duplex determines the site of template switching

To determine the effect of the TRS duplex formation upon the incidence of template switching, we modelled the RNA structures formed between cTRS-L and TRS-B and surrounding sequences and compared these with junctional sequences and the abundance of transcripts determined by Kim et al. (2020). The abundance is a measure of the steady state levels of the RNA species rather than a direct measure of nascent transcript production. Therefore, the abundance would represent a minimum steady state level of the sgRNA species. We modelled duplex structures of the 5’ leader sequence (up to 94 nts) which includes SL1-SL3 and a small section of SL4, and the nascent negative strand sequence with varying lengths of nucleotides before and after the cCS-B, beyond the site of complementarity, and compared these to the transcript abundance. We do not show structures associated with TRS-L/cTRS-B-independent fusion events. We also modelled body sequences of different lengths, up to 6 nucleotides beyond the cCS, to simulate the structures that may be formed when the polymerase has produced a longer nascent negative strand. In some cases, changing the length of the nucleotide sequence modifies the RNA secondary structure. In these cases, we have depicted them separately.

By comparing the duplex structures to junctional sequences between the 5’ leader and the cTRSs in the body of the nascent negative strand, we observe that in the most abundant sgRNAs, a majority of template switching appears to occur within or immediately after the last base-paired nucleotide in the CS-L:cCS-B duplex (i.e. 5’ of the duplex on the +strand) or 1-2 nucleotides after the duplex (less commonly). Due to the shared core sequence at the CS-L and the CS-B, in most cases, the junctional sequence products of template switching events that have occurred within the duplex or immediately after, look identical. All of these events have been depicted with a single arrow at the last possible site of template switching. Template switching was also found in bulges or at weak G-U base pairs. This is seen for the most abundant sgRNAs for S, ORF3a, E, M, ORF6, ORF7a, ORF8, and N (Figs. 2, 3, 4). All the duplexes shown are more stable (lower Gibbs free energy) than the structure where the template switch-causing duplex is not formed. The exception is the ORF6 duplex which showed the same free energy of secondary structure formation for the structure with the duplex as the structure without the template switch-causing duplex where the cCS-B is sequestered in a weak stem-loop (Fig. 2). This may explain why this sgRNA is the 8^th^ most abundant sgRNA.

**FIGURE 2.**
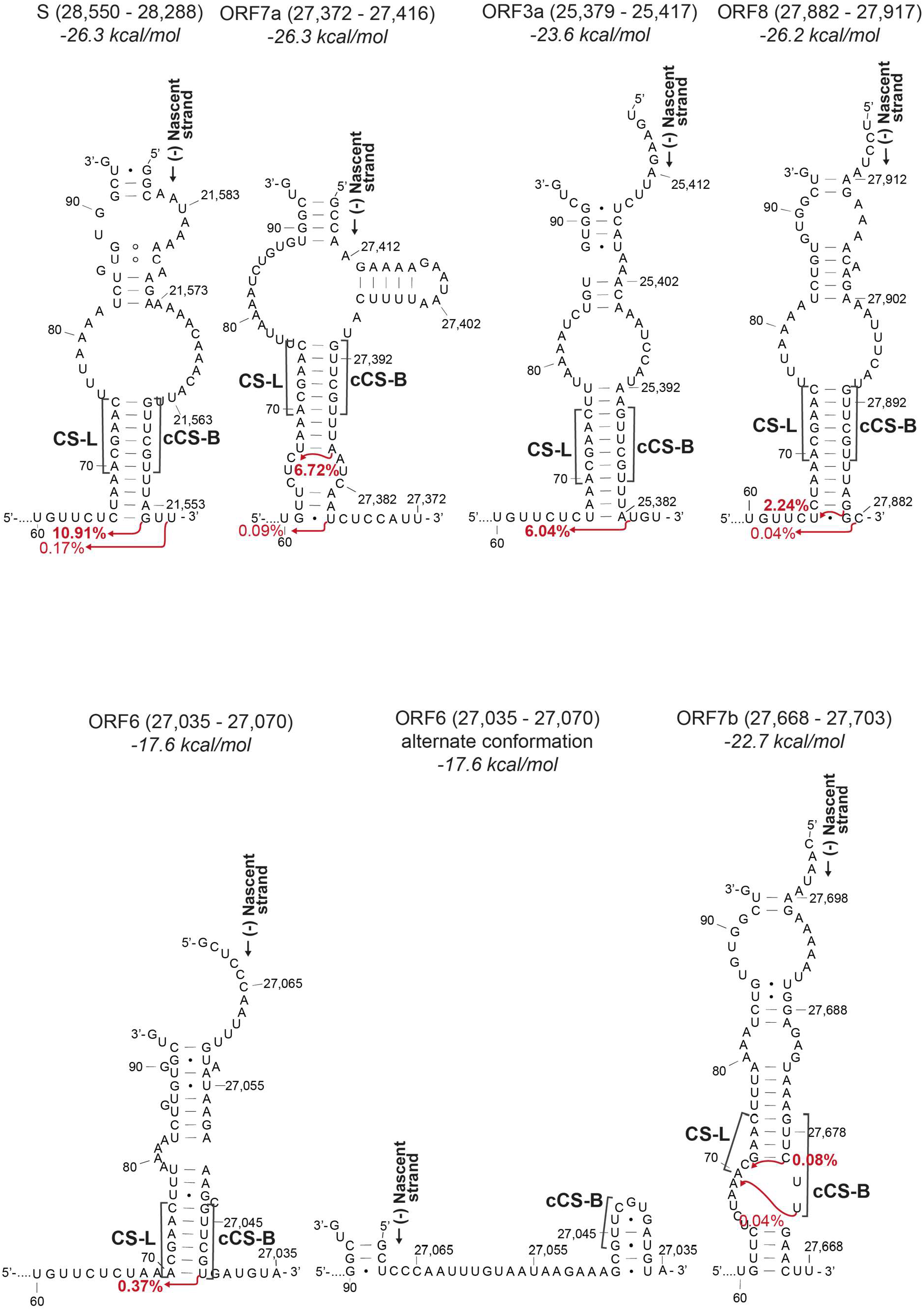
Duplexes formed between the nascent transcript and the 5’ UTR on the positive strand leading to sgRNAs S, ORF7a, ORF3a, ORF8, ORF6 and ORF7b. The start of the 5’ leader sequence and the nascent strand are indicated. The red arrows indicate the position after which template switching occurs and the percentage of total reads associated with the resultant leader-body junction (data from Kim et al. (2020)). The nucleotide numbering indicates the position in the positive strand. The labels above indicate the region of the nascent strand depicted and the folding free energy for each of the models. For the second and third structure, the 5’ UTR up to U60 was also modelled but is not shown. CS-L – proposed leader core sequence in the 5’ UTR, cCS-B – Sequence complementary to the core sequence in the body

**FIGURE 3.**
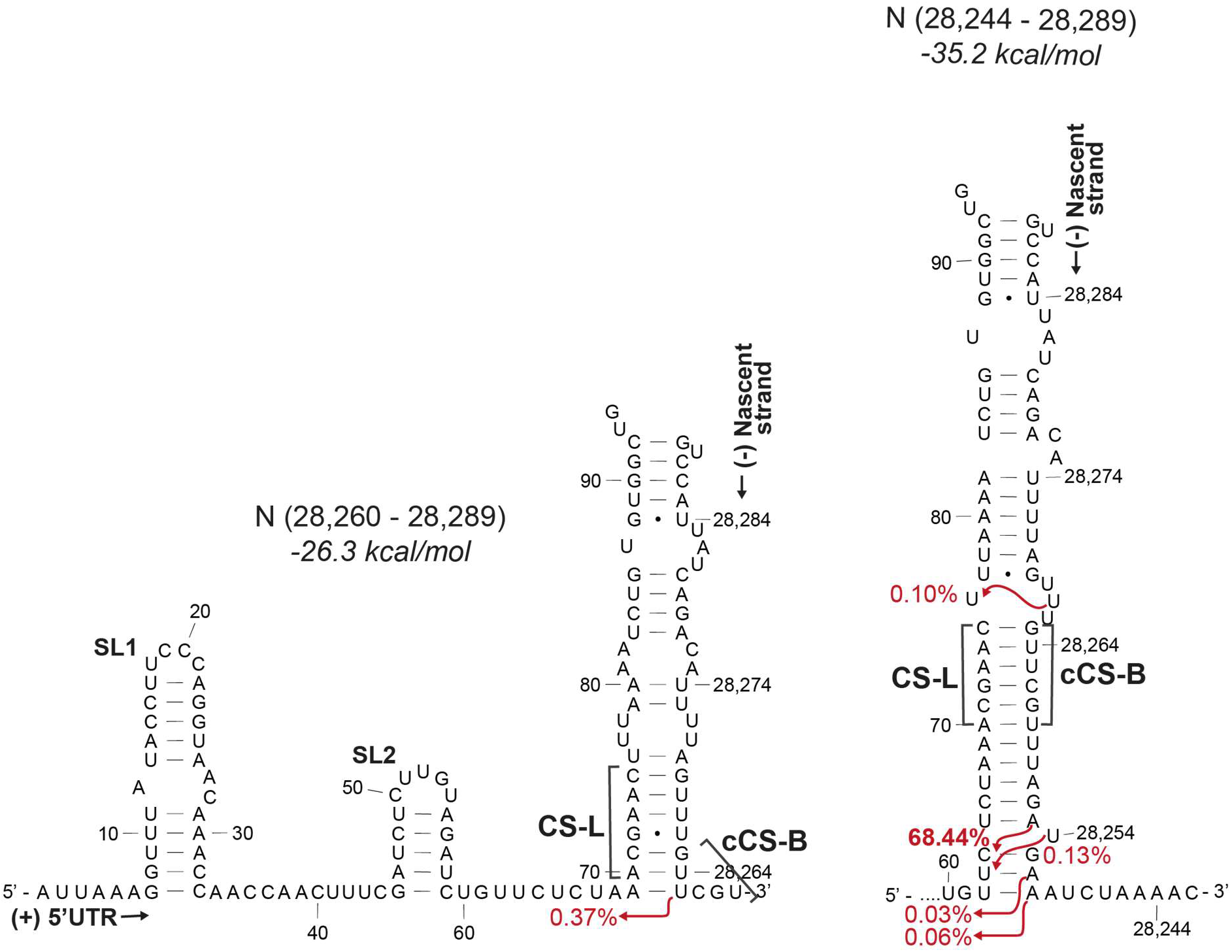
Duplexes formed between the nascent transcript at the N TRS-B and the 5’ UTR. The second most abundant sgRNA is formed from a non-canonical duplex with a sequence adjacent to the core sequence (left-hand side). CS-L – proposed leader core sequence in the 5’ UTR, cCS-B – Sequence complementary to the core sequence in the body, SL1 – Stem-loop 1, SL2 – Stem-loop 2

**FIGURE 4.**
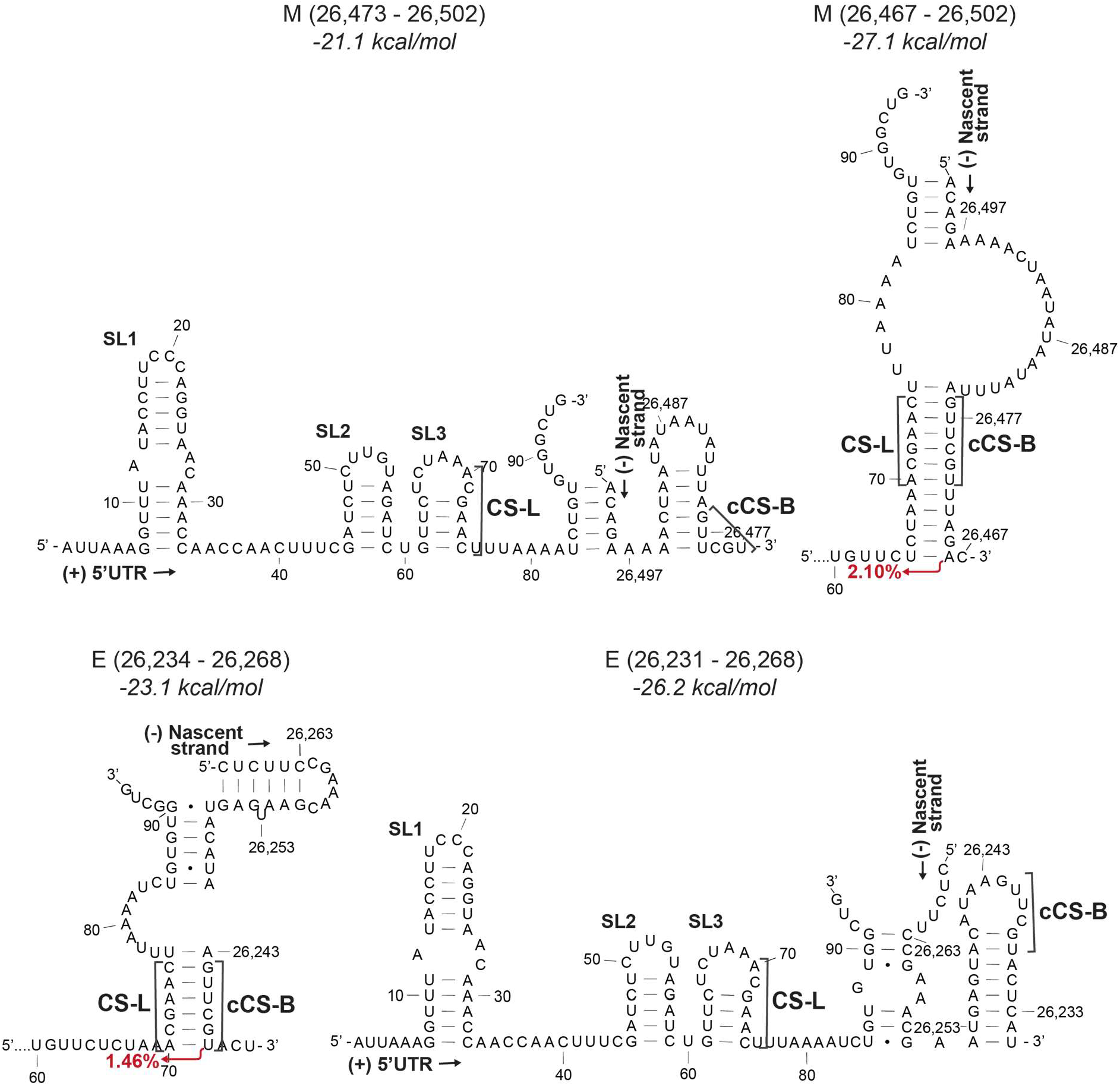
Duplexes formed between the 5’ UTR and nascent transcript at the M TRS-B (above) or the E TRS-B (below) of different lengths. The nascent strands are shorter on the left. The free energy for the secondary structure is given above. CS-L – proposed leader core sequence in the 5’ UTR, cCS-B – Sequence complementary to the core sequence in the body, SL1 – Stem-loop 1, SL2 – Stem-loop 2, SL3 – Stem-loop 3

The second most abundant ORF7a sgRNA which accounts for 0.09% of total transcripts is produced from a template switching event occurring after a bulge and a second duplex beyond the primary CS-L:cCS-B duplex (Fig. 2). This is also seen in the lower abundance N sgRNAs (0.03% and 0.06% abundance) where template switching occurs after the site of a one-nucleotide bulge (Fig. 3). It may be that in some cases a small bulge is disregarded during template switching if it is followed by another duplex, perhaps indicating the coaxial stacking of those helices on either side of a single nucleotide bulge ^30^.

### Speed of transcription affects duplex formation

Of the 23 sgRNAs studied, six of them are N sgRNAs (68.05%, 0.37%, 0.13%, 0.10%, 0.06% and 0.03% abundance). This may be due to its 3’ terminal position in the genome making it the first TRS-B sequence that the polymerase encounters or may be a function of sequence composition or both. The second most abundant alternate N sgRNA (0.37%) is not formed from the canonical CS-L:cCS-B duplex (Fig. 3, left hand panel). The TRS-B for the N sgRNA is found within a 5’ACNAACNAAC3’ sequence where the CS-L can base pair with the complement of both ACNAAC sections. Topologically, the cCS-B is the second CS to be produced in the nascent negative strand. However, unlike the alternate cCS-B which relies on a G-U base pairing for duplex formation, the duplex formed with a longer nascent -ve strand and the canonical cCS-B is predicted to be more stable and thus leads to a higher incidence of template switching (Fig. 3, right hand panel). The availability of multiple template switching points, all producing the same N ORF, indicates a viral strategy favouring N sgRNA synthesis and contributing consequently to making the nucleocapsid protein the most abundantly expressed protein in infected cells ^31^.

The length of the nascent negative transcript is closely related to the speed of polymerase progression. A slower polymerase would allow more time for a duplex to be produced earlier in the nascent transcript. AU-rich regions have been previously described to promote more efficient recombination and template switching than GC-rich sequences in unrelated RNA dependent RNA polymerases in plant viruses ^32^. One of the SARS-CoV-2 N sgRNAs (0.10%) is unusual in that it is created from a template switching event that occurs even before the core sequence (Fig. 3, right hand panel). This occurs after a stretch of As and Us, homopolymeric runs of which are known to be associated with frequent polymerase pausing and dissociation and thus will favour template switching events ^33^.

The effect of the polymerase processivity and the length of the nascent strand on template switching is also evident in simulations of the formation of the M and E sgRNAs. If a nascent transcript is produced that ends with the M cCS-B, the cCS-B is partly sequestered in a stem-loop that is more stable than a duplex with the CS-L. However, if a longer transcript is produced, a duplex is formed between the CS-L and cCS-B which leads to template switching (Fig. 4A). Conversely, a shorter nascent transcript that ends closer to the cCS-B in the E sgRNA is more likely to lead to the CS-L:cCS-B duplex that results in template switching, than a longer transcript where the cCS-B is partly sequestered in a stem-loop. This sequestration of the cCS-B in the stem-loop may also prevent the inhibition of downstream duplex formation.

### The non-canonical ORF7b CS-B prevents frequent template switching

Of the four ORF7b sgRNAs shown, the sgRNAs are predicted to produce 20 (0.08%), 4 (0.06%), 43 (0.04%) and 43 (0.02%) amino acid peptides (in order of decreasing transcript abundance). The least abundant 43 aa ORF7b-producing sgRNA (0.02%) is not produced from a CS-L:cCS-B mediated fusion event, but rather from within the body of the genome further upstream from the ORF7b CS-B. It is not yet clear if these RNAs are translated as the presence of the two shorter translation products has not been experimentally confirmed.

Of the 24 most abundant RNAs produced from the viral genome, only two sgRNAs (with a total abundance of 0.056%) encode the full ORF7b 43 aa protein. Of all the RNA species detected in the Kim et al. (2020) paper, 0.085% encode the full protein. The low abundance of these RNA species is caused by a proposed non-canonical CS-B 5’AACAAG3’ which does not produce a complement that is able to form a perfect duplex with the 5’ACGAAC3’ CS-L. When recombination does occur the most common site of template switching within the CS-L, while the other frequent event occurs immediately after the cCS-B. However, in both cases, template switching still occurs immediately after one or to two nucleotides after a duplex, as is observed with the other sgRNAs. The low levels of ORF7b coding sgRNAs may have evolved to reduce dosage of the protein. Interestingly, in SARS-CoV, ORF7b, which may be a viral structural protein, is translated by an inefficient mechanism of leaky ribosomal scanning possibly with the same consequence of limiting its quantity ^34^.

## DISCUSSION

The abundance of sgRNA species and thus the translational expression of the virally encoded proteins is controlled by TRS-mediated duplex formation. Structural modelling of the duplex formed between the SARS-CoV-2 negative strand nascent transcript and the positive strand leader sequence identified stable structures. Here we show that the site of template switching correlates with the length and stability of the duplex. In most cases, the polymerase probably switches templates from within the CS-L:cCS-B duplex up to 1-2 nucleotides after the duplex. Since the chimeric RNA created from a template switching event that occurs within the complementary duplex sequence would look identical regardless of where the template switching event occurs within the duplex, it would be difficult to conclude that all the modelled template switching events happen after the duplex. Indeed, 0.10% of all sgRNAs are produced from template switching that occurs even before the CS-L:cCS-B duplex in the case for the second most abundant N sgRNA. In the case of the alternate N sgRNA that is produced from a non-canonical duplex, the presence of a G-U base pair in the duplex indicates that the template switching event must occur after this point (Fig 3). Since different lengths of nascent RNAs can sequester the cTRS-B in a structure, thus preventing contact between the leader and body sequences, a minimum length and stability of duplex dependent on the sequence composition would be required.

In the case of the N sgRNA, multiple highly abundant RNA products are produced from different template switching events (Fig 3). This may be due to its proximity to the 3’ end of the genome. However, this positional argument is not substantiated by the high abundance of other sgRNAs which are not also relatively 3’ proximal. Alternatively, the presence of an alternate duplex conformation increases the likelihood of template switching and nucleocapsid production. The high abundance of nucleocapsid protein enables multiple RNA contact points, which may not only protect the RNA from innate immune recognition, but also encourage oligomerisation and genome encapsidation ^35,36^. This is caused by the recruitment of DDX1 helicase mediated by N that enables progression of the polymerase to the 5’ end of the genome ^37^. Thus, increased nucleocapsid production would not only be involved in a positive feedback loop regulating discontinuous transcription and further nucleocapsid production, but also, as a packaging protein, this could allow temporal control of when the gRNA is prioritised for packaging over transcription. Contrasting with this, the absence of a canonical core sequence at the 5’ terminus of ORF7b prevents the formation of strong duplexes and thus produces low levels of ORF7b encoding sgRNA. Alternatively, more abundant ORF7b sgRNAs produced at the associated duplex could produce other as yet unidentified translation products from alternate ORFs ^3^.

Template switching seems to be controlled by the length of the nascent negative transcript. This could be due to the sequestration of the cTRS-B in their own stem-loops. Structure modelling of the positive strand also revealed that most TRS-B sequences are found in secondary structures ^20^. Recent DMS-MaPseq of the SARS-CoV-2 positive strand RNA in the cell showed that, except for the TRS-B for ORF7a and ORF7b, all the TRS-Bs were found completely or partially enclosed in secondary structures ^38^. Four of these TRSs were found to be highly stable ^21^. From the modelling data presented here, we see the formation of cTRS-B stem-loops in the negative strand which reflect the structures observed in the positive strand (Fig 4).

We posit that such a mechanism has three roles. Firstly, the melting of the structures required for RNA synthesis slows down the approaching polymerase on the template. It has been shown that structures and composition of the template, such as the presence of homopolymeric runs of nucleotides, particularly adenosines, sequences as seen in the N duplex, can cause pausing of polymerases ^33^. Then, the deceleration or dissociation/reattachment of the polymerase would allow further opportunities for duplex formation between the cTRS-B and TRS-L.

Secondly, the formation of the structures prevents the inhibition of downstream TRS-Bs. It has previously been reported that, in the event of two closely positioned TRSs, the downstream TRS may inhibit the upstream TRS but not vice versa ^39,40^. We postulate that the reformation of secondary structures in the nascent negative strand prevents inappropriate TRS-L:cTRS-B duplex formation and allows the polymerase to progress to the site of the next TRS-B. This process of unwinding and reformation of secondary structures before and after the polymerase would be vital again during RNA-dependent RNA synthesis of the positive strand.

Thirdly, the chances of triggering the cellular dsRNA detection mechanisms are reduced. Virally encoded nsp13 helicase unwinds the dsRNA duplexes produced during replication or transcription ^41^. We speculate that the formation of secondary structures on the unwound nascent negative strand in cis may prevent the reformation of long dsRNA between the template and the nascent strand and may thus provide an additional mechanism by which the virus avoids dsRNA-triggered host cell responses.

We conclude that the strength and stability of secondary structures during nascent RNA negative strand synthesis at the site of the TRS-B provides a plausible explanation for regulation of numerous mechanisms involved in discontinuous transcription including the site of template switching. Experimental studies will be required to confirm these conclusions. Interfering with structures formed during discontinuous transcription may serve as potential RNA targets for therapeutic intervention.

## ACKNOWLEDGEMENTS

Dr Ulrich Desselberger is acknowledged for very helpful discussions.

The work was funded by the Clinical Academic Reserve (Cambridge), the Biomedical Research Centre (Cambridge) and grants from NUS Department of Medicine

## CONFLICT OF INTEREST STATEMENT

The authors declare that they have no conflicts of interest

## Notes

### Competing Interest Statement

The authors have declared no competing interest.

